# Promoter or enhancer activation by CRISPRa rescues haploinsufficiency caused obesity

**DOI:** 10.1101/140426

**Authors:** Navneet Matharu, Sawitree Rattanasopha, Lenka Maliskova, Yi Wang, Aaron Hardin, Christian Vaisse, Nadav Ahituv

## Abstract

Haploinsufficiency, having only one functional copy of a gene, leads to a wide range of human disease and has been associated with over 300 genes. Here, we tested whether CRISPR activation (CRISPRa) could rescue a haploinsufficient disease *in vivo*. Haploinsufficiency of *Sim1*, a transcription factor involved in the leptin pathway, results in severe obesity in humans and mice. CRISPRa targeting of either the *Sim1* promoter or its ~270kb distant hypothalamic enhancer using transgenic mice, rescued the obesity phenotype in *Sim1* heterozygous mice. Interestingly, despite using a ubiquitous promoter for CRISPRa, *Sim1* was upregulated only in tissues where the promoter or enhancer are active, suggesting that cis-regulatory elements can determine CRISPRa tissue-specificity. To further relate this to therapy, we injected CRISPRa adeno associated virus into the hypothalamus, leading to reversal of the obesity phenotype. This therapeutic strategy could be used to rescue numerous diseases resulting from altered gene dosage.

## Introduction

Over 300 genes are known to cause human disease due to haploinsufficiency^1,2^, leading to a wide range of phenotypes that include cancer, neurological diseases, developmental disorders, immunological diseases, metabolic disorders, infertility, kidney disease, limb malformations and many others^1^. Large-scale exome sequencing analyses estimate that a total of 3,230 human genes could be heterozygous loss-of-function (LoF) intolerant^3^. Gene therapy holds great promise in correcting haploinsufficient diseases, by inserting a functional recombinant copy or copies of the mutant gene. Currently, there are a total of 2,300 clinical trials underway for gene therapy, the majority of them using adeno-associated virus (AAV) to deliver the recombinant gene^4^. AAV is a preferred gene delivery method due to its ability to deliver DNA without integrating into the genome, not causing pathogenicity and providing long lasting gene expression of the transgene^5^. However, current AAV approaches tend to use ubiquitous promoters to drive transgenes that can lead to non-desirable ectopic expression. Another crucial limitation is that AAV has an optimal 4.7 kilo base (kb) packaging capacity, limiting its gene therapy use for genes longer than 3.5kb (taking into account additional regulatory sequences needed for its stable expression). Analysis of the 300 haploinsufficiency disease causing genes and 3,230 predicted heterozygous LoF genes finds 156 (52%) and 715 (22%) of them, respectively, to have coding sequences longer than 3.5kb (**Extended Data Fig. 1**), rendering them not suitable for AAV gene therapy. CRISPR gene editing can potentially fix haploinsufficient mutations, however it would require the need to custom tailor the editing strategy for each mutation. Moreover, it’s not a feasible therapy for heterozygous LoF micro-deletions. To address these challenges, we devised a novel therapeutic strategy to treat haploinsufficiency using CRISPR activation (CRISPRa). CRISPRa takes advantage of the RNA-guided targeting ability of CRISPR to direct a nuclease deficient Cas9 (*dCas9*) along with a transcriptional activator to regulatory element/s of a specific gene, thus increasing its expression^6–10^. Here, we tested whether we can use this system to increase the transcription of the unaffected endogenous gene in a haploinsufficient disease to rescue the disease phenotype. As a proof-of-concept model, we chose a quantitative trait, obesity caused by haploinsufficiency of the single-minded family bHLH transcription factor 1 (*SIM1*) gene.

*SIM1* gene is a transcription factor that is expressed in the developing kidney and central nervous system, and is essential for the formation of the supraoptic nuclei (SON) and paraventricular nuclei (PVN) of the hypothalamus^11^. It is also thought to play a major role in the leptin pathway^12^. In humans, haploinsufficiency of *SIM1* due to chromosomal aberrations results in hyperphagic obesity^13^ and *SIM1* coding mutations, many of them being loss-of-function, are thought to be a major cause of severe obesity in humans^14–16^. *Sim1* homozygous null mice die perinatally, while *Sim1* heterozygous mice (*Sim1*^+/−^) survive, are hyperphagic and develop early-onset obesity with increased linear growth, hyperinsulinemia and hyperleptinemia^17^. A postnatal conditional knockout of hypothalamic *Sim1* leads to a similar phenotype in heterozygous mice^18^, delineating an additional role for *Sim1* as an important regulator of energy homeostasis in adults. Overexpression of *SIM1*, using a human bacterial artificial chromosome in mice, rescues diet-induced obesity and reduced food intake^19^, suggesting a potential role for *Sim1* as a therapeutic target for obesity. We tested the ability of CRISPRa to rescue the obesity phenotype in *Sim1*^+/−^ mice using both transgenic and AAV based approaches. CRISPRa using an sgRNA targeted to either the *Sim1* promoter or its ~270 kb distant enhancer upregulated its expression and rescued *Sim1* mediated obesity in transgenic animals. AAV-mediated delivery of CRISPRa to the hypothalamus prevented excessive weight gain in postnatal *Sim1*^+/−^ mice. Our results present a novel therapeutic approach for treating haploinsufficient disease.

## Results

### Upregulation of *Sim1 in vitro*

We first set out to optimize our CRISPRa conditions *in vitro. Sim1* has a well characterized promoter^20^ and distant and robust hypothalamus enhancer (~270kb from the transcription start site), *Sim1* candidate enhancer 2 (SCE2^21^), both of which were chosen as targets for CRISPRa (**Fig. 1a**). We designed two sgRNAs for either the *Sim1* promoter or SCE2. Using these guides we tested if dCas9 fused to VP64 (dCas9-VP64), a transcriptional activator that carries four tandem copies of VP16 (a herpes simplex virus type 1 transcription factor)^22^, can upregulate *Sim1* in mouse neuroblastoma cells (Neuro-2a). The VP64 activator domain was chosen due to its small size and moderate activation potential compared to other known activators^23^, as we wanted to obtain therapeutic *Sim1* dosage levels *in vivo* that are similar to wild-type. Cells were transfected with dCas9-VP64 and the various guides and following 48 hours *Sim1* mRNA levels were measured using quantitative PCR (qPCR). We identified one sgRNA for either promoter or SCE2 that was able to upregulate endogenous *Sim1* by 13 and 4 fold respectively (**Fig. 1b; Extended Data Fig. 2**).

**Figure 1.**
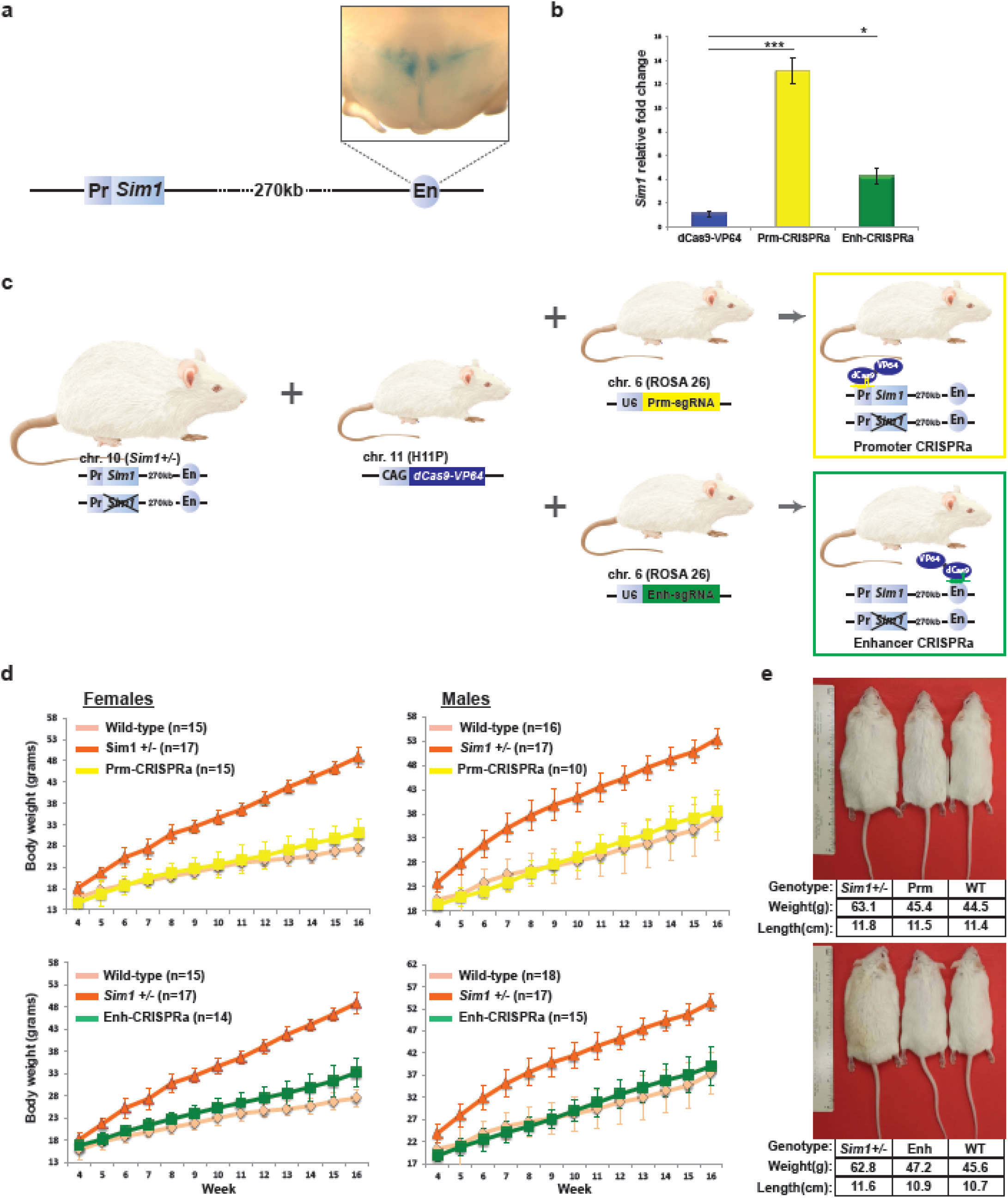
CRISPRa *Sim1* upregulation *in vitro* and obesity rescue *in vivo*. **a**, Schema of the mouse *Sim1* genomic locus, showing the LacZ driven hypothalamus expression of SCE2 (En) from 56 day old mice. **b**, CRISPRa in Neuro-2A cells targeting the *Sim1* promoter (Prm-CRISPRa) or enhancer (Enh-CRISPRa). Results are expressed as mRNA fold-increase normalized to beta-actin using the ΔΔCT method. The mean values±s.d. were obtained from 3 independent experiments. * = p-value < 0.001 *** = p-value < 0.0005 (ANOVA, Tukey test). c, Schema showing the mating scheme used to generate *Sim1*^+/−^ CRISPRa mice. A CAG-dCas9-VP64 cassette was knocked in *Hipp11* (H11P) locus and an sgRNA targeting either the *Sim1* promoter (U6-Prm-sgRNA) or sCe2 (U6-Enh-sgRNA) were knocked into the *Rosa26* locus.d, Weekly weight measurements of wild-type, *Sim1*^+/−^, *H11P^pAG-dCas9-VP64^* X *ROSA26^Sim1Pr-sgRNA^* (Prm-CRISPRa) and *H11P^CAG-dCas9-VP64^* X *ROSA26^SCE2En-sgRNA^* (Enh-CRISPRa). At least 10 male and female mice were measured per genotype. Mean values±s.d are shown. e-f, Pictures showing 26 week old male mice for each genotype: *Sim1*^+/−^, *H11P^CAG-dCas9-VP64^* X *ROSA26^Sim1Pr-sgRNA^* (Prm) and wild-type (WT) (e) and *Sim1*^+/−^, *H11P^CAG-dCas9-VP64^* X *ROSA26^SCE2En-sgRNA^* (Enh) and wild-type (WT) (f). Genotype, weight and length of each mouse are depicted below.

### Transgenic CRISPRa rescues obesity

To test the ability of our CRISPRa system to rescue *Sim1* obesity *in vivo*, we generated knockin mouse lines using TARGATT technology^24^ that have dCas9-VP64 inserted into the mouse *Hipp11* (*H11p^CAG-dCas9-VP64^*) locus and either sgRNA, targeting the *Sim1* promoter (*ROSA26^Sim1Pr-sgRNA^*) or SCE2 (*ROSA26^SCE2En-sgRNA^*), in the *Rosa26* locus (**Fig. 1c, Extended Data Fig. 3**). We then crossed these mice to *Sim1*^+/−^ mice that develop severe obesity^17^. Mice having all three alleles (*Sim1*^+/−^ X *H11P^CAG-dCas9-VP64^* and *ROSA26^Sim1Pr-sgRNA^* or *ROSA26^SCE2En-sgRNA^*) were maintained using breeders chow (picodiet-5058) and weighed on a weekly basis until 16 weeks of age along with wild-type littermates and *Sim1*^+/−^ and *Sim1*^+/−^ X *H11P^CAG-dCas9-VP64^* mice both of which become severely obese (negative controls). Analysis of at least ten females and ten males per condition showed that *Sim1*^+/−^ mice carrying both dCas9-VP64 and either *Sim1* promoter or enhancer sgRNA have a significant reduction in body weight compared to *Sim1*^+/−^ X *H11P^CAG^’^dCas9^’^VP64^* and *Sim1*^+/−^ (**Fig. 1d-e; Extended Data Fig 4**). Wild-type mice carrying both dCas9-VP64 and either *Sim1* promoter or enhancer sgRNA also showed reduction in body weight compared to wild-type mice (**Extended Data Fig 4**).

To relate body weight reduction with body composition and metabolic parameters, we next performed metabolic profiling for *Sim1*^+/−^ X *H11P^CAG-dCas9-VP64^* X *ROSA26^Sim1Pr-sgRNA^* (Prm CRISPRa) *Sim1*^+/−^ X *H11P^CAG-dCas9-VP64^* X *ROSA26^SCE2En-sgRNA^* (Enh-CRISPRa) *Sim1*^+/−^ and wild-type mice. Three male and three female mice for each genotype were analyzed for body composition and metabolic profiling, during the onset of the obesity phase, 8 weeks of age. Both Prm-CRISPRa and Enh-CRISPRa mice showed significantly reduced body fat content compared to *Sim1*^+/−^ in both females and males (**Fig. 2a**). Metabolic chamber analyses of other hallmarks of *Sim1*^+/−^ obese mice, such as oxygen consumption and food intake, showed a shift towards wild-type metabolic parameters in the Prm-CRISPRa and Enh-CRISPRa mice (**Fig. 2b-c**). In addition, their respiratory exchange ratio (RER; VCO2/VO2), an indirect method of defining basic metabolic rate, also showed parameters similar to their wild-type littermates (Fig. 2d). We did not observe any significant physical activity differences in individual chambers (Extended Data Fig. 5). Combined, these results show that both Prm-CRISPRa and Enh-CRISPRa mice have reduced body weight due to lower food intake, which likely leads to their reduced body fat levels and an overall improvement in their metabolic parameters.

**Figure 2.**
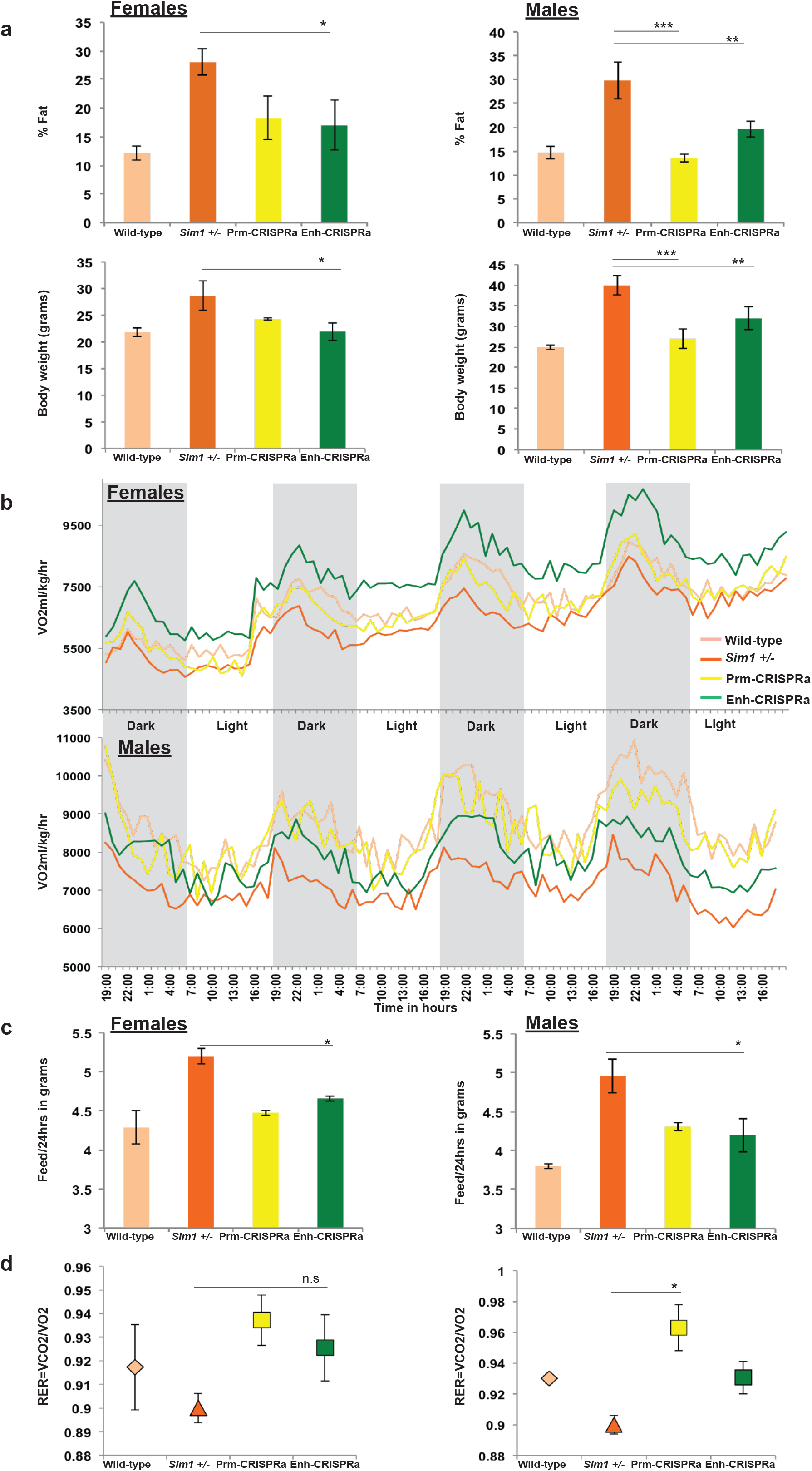
Body composition and metabolic analyses of *Sim1* CRISPRa transgenic mice. **a**, Measured percent fat in wild-type littermates, *Sim1*^+/−^, *H11P^CAG-dCas9-VP64^*X *ROSA26^Sim1Pr-sgRNA^* (PrmCRISPRa) and *H11P^CAG-dCas9-VP64^*X *ROSA26^SCE2En-sgRNA^* (EnhCRISPRa) as determined by Echo Magnetic Resonance Imaging (EchoMRI) for females and Dual Energy X-ray Absorptiometry (DEXA) for males, with their corresponding body weight measurements. The mean values±s.d. were obtained from 3 females and 3 males. **b**, Metabolic chamber VmaxO2 measurements averaged hourly for 3 males and 3 females for all four genotypes: wild-type, *Sim1*+/−, Prm-CRISPRa and Enh-CRISPRa determined over a 4 day period. **c**, Food intake per day for all four genotypes determined over a 4 day period. Mean values±s.d. were obtained from 3 females and 3 males. * = p-value < 0.01; *** = p-value < 0.005; n.s = non-significant (ANOVA, Tukey test). **d**, Respiratory exchange ratio (RER; VCO2/VO2) for all four genotypes obtained from 3 females and 3 males and plotted as mean values±s.d.

### *Sim1* activation is tissue-specific

To test for *Sim1* activation levels and tissue-specificity in Prm-CRISPRa and Enh-CRISPRa mice, we measured its mRNA expression levels in different tissues. We selected two tissues where *Sim1* is known to be expressed, hypothalamus and kidney, and two tissues where it is not expressed, lung and liver, based on previous studies^25,26^ and our analysis of *Sim1* expression in different tissues (**Extended Data Fig. 6**). We first measured *dCas9* expression, and found it to be expressed in all four tissues, as expected, since we used a ubiquitous CMV enhancer chicken beta-Actin (CAG) promoter to drive its expression (**Fig. 3a**). In contrast, for *Sim1*, we observed significantly higher mRNA levels in both the hypothalamus and kidney in Prm-CRISPRa mice but only in the hypothalamus of Enh-CRISPRa mice compared to *Sim1*^+/−^ mice (**Fig. 3b**). In *Sim1*^+/−^ mice, as expected, we observed half the levels of mRNA when compared to wild-type mice both in the hypothalamus and kidney. To check if *Sim1* promoter or enhancer CRISPRa has an effect on neighboring genes, we analyzed the mRNA expression levels of activating signal cointegrator 1 complex subunit 3 (*Ascc3*) and G protein-coupled receptor class C group 6 member A (*Gprc6a*). We did not observe any differences in expression levels for these genes in Prm-CRISPRa and Enh-CRISPRa compared to wild-type mice (**Extended Data Fig. 7a-b**). To assess targeting specificity, we carried out chromatin immunoprecipitation (ChIP) for *dCas9* followed by qPCR for *Ascc3, Gprc6a* and *Sim1* promoters or SCE2. We observed *dCas9* binding only in the *Sim1* promoter for Prm-CRISPRa and SCE2 in Enh-CRISPRa mice (**Extended Data Fig. 7c-d**).

**Figure 3.**
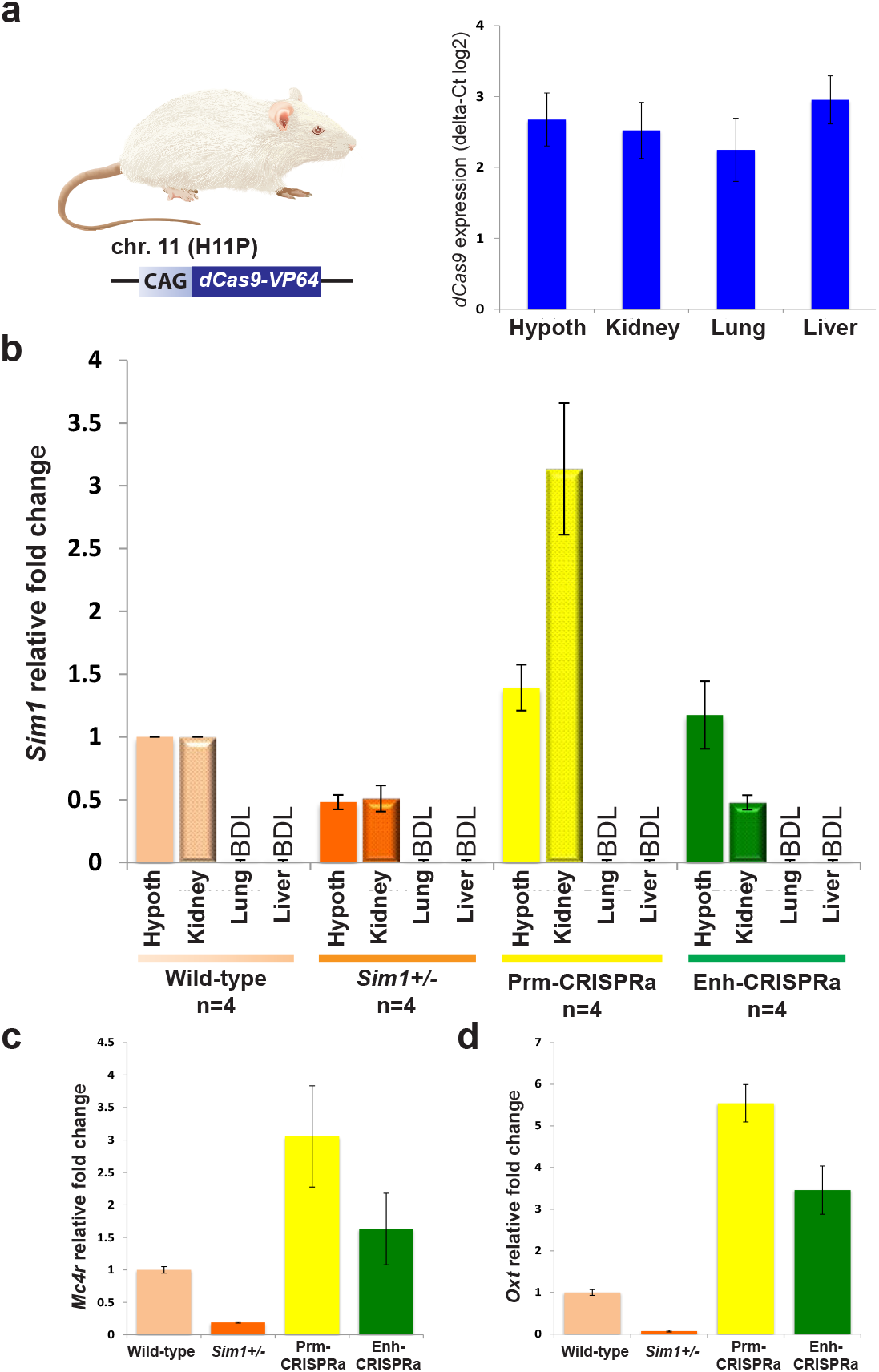
*dCas9* and *Sim1* mRNA expression levels in CRISPRa transgenic mice. **a**, *dCas9* mRNA expression in the hypothalamus, kidney, lung and liver from four *Sim1*^+/−^ X *H11P^CAG-dCas9-VP64^* mice. **b**, *Sim1* mRNA expression in the hypothalamus, kidney, lung and liver for the following genotypes: wild-type, *Sim1*^+/−^, *H11P^CAG-dCas9-VP64^* X *ROSA26^Sim1Pr-sgRNA^* (Prm-CRISPRa) and *H11P^CAG-dCas9-VP64^* X *ROSA26^SCE2En-sgRNA^* (Enh-CRISPRa) from 4 mice (2 females and 2 males). **c-d**, *Mc4r* (**c**) and *Oxt* (**d**) mRNA expression levels in the hypothalamus for the following genotypes: wild-type, *Sim1*^+/−^, Prm-CRISPRa and Enh-CRISPRa from 4 mice (2 females and 2 males). The mean values±s.d for all experiments were determined based on mRNA fold-increase compared to wild-type and normalized to *beta-actin* or *Rpl38* using the ΔΔCT method or ΔCT for **a**. BDL = below detectable levels.

Since we did not observe any significant differences between the obesity phenotype of Prm-CRISPRa and Enh-CRISPRa mice, we could speculate that the activation of *Sim1* in the hypothalamus is sufficient to rescue the *Sim1*^+/−^ obesity phenotype. Supporting our results a previous hypothalamus specific postnatal conditional knockout of *Sim1* was also shown to cause obesity in heterozygous mice and reduction in melanocortin 4 receptor (*Mc4r*) and oxytocin (*Oxt*) mRNA levels^18^, both downstream factors in the leptin pathway. Analysis of *Mc4r* and *Oxt* mRNA levels in the hypothalamus showed an increase in their expression in Prm-CRISPRa and Enh-CRISPRa mice (**Fig. 3c-d**), suggesting that *Sim1* upregulation can lead to their increased expression. Interestingly, in tissues where *Sim1* is not expressed (*i.e*. liver and lung), we could not detect *Sim1* expression in Prm-CRISPRa or Enh-CRISPRa mice despite *dCas9* being expressed. These results imply that in spite of ubiquitous expression, dCas9-VP64 could only upregulate *Sim1* in tissues where its target *cis*-regulatory elements are active. This suggests that cis-regulatory elements could be used to define the tissue-specificity of CRISPRa.

### CRISPRa AAV decreases *Sim1*^+/−^ weight gain

To further translate this approach to a therapeutic strategy for haploinsufficiency, we took advantage of AAV to deliver CRISPRa into the hypothalamus of *Sim1*^+/−^ mice. We generated the following three AAV vectors: 1) dCas9-VP64 driven by a cytomegalovirus (CMV) promoter (*pCMV-dCas9-VP64*); 2) *Sim1* promoter sgRNA along with mCherry (*pU6-Sim1Pr-CMV-mCherry*); 3) SCE2 sgRNA along with mCherry (*pU6-SCE2-CMV-mCherry*) (**Fig. 4a**). For the *pCMV-dCas9-VP64* vector, due to the size of dCas9-VP64 expression cassette, we obtained a 5.4kb insert. While this insert size is above the 4.7kb optimal AAV packaging limit, it was shown that going above 5kb reduces transgene expression levels but still could be used for delivery^27^. These vectors were packaged individually into AAV-DJ serotype, which is a chimera of type 2, 8 and 9 that was shown to achieve high expression levels in multiple tissues^28^. We did observe lower but usable viral titers for *pCMV-dCas9-VP64* AAV (see methods). We first tested if our AAV CRISPRa vectors could upregulate *Sim1 in vitro* using Neuro-2a cells. We observed a 4 and 5 fold upregulation of *Sim1* mRNA expression when targeting the promoter or enhancer respectively (**Fig. 4b**).

**Figure 4.**
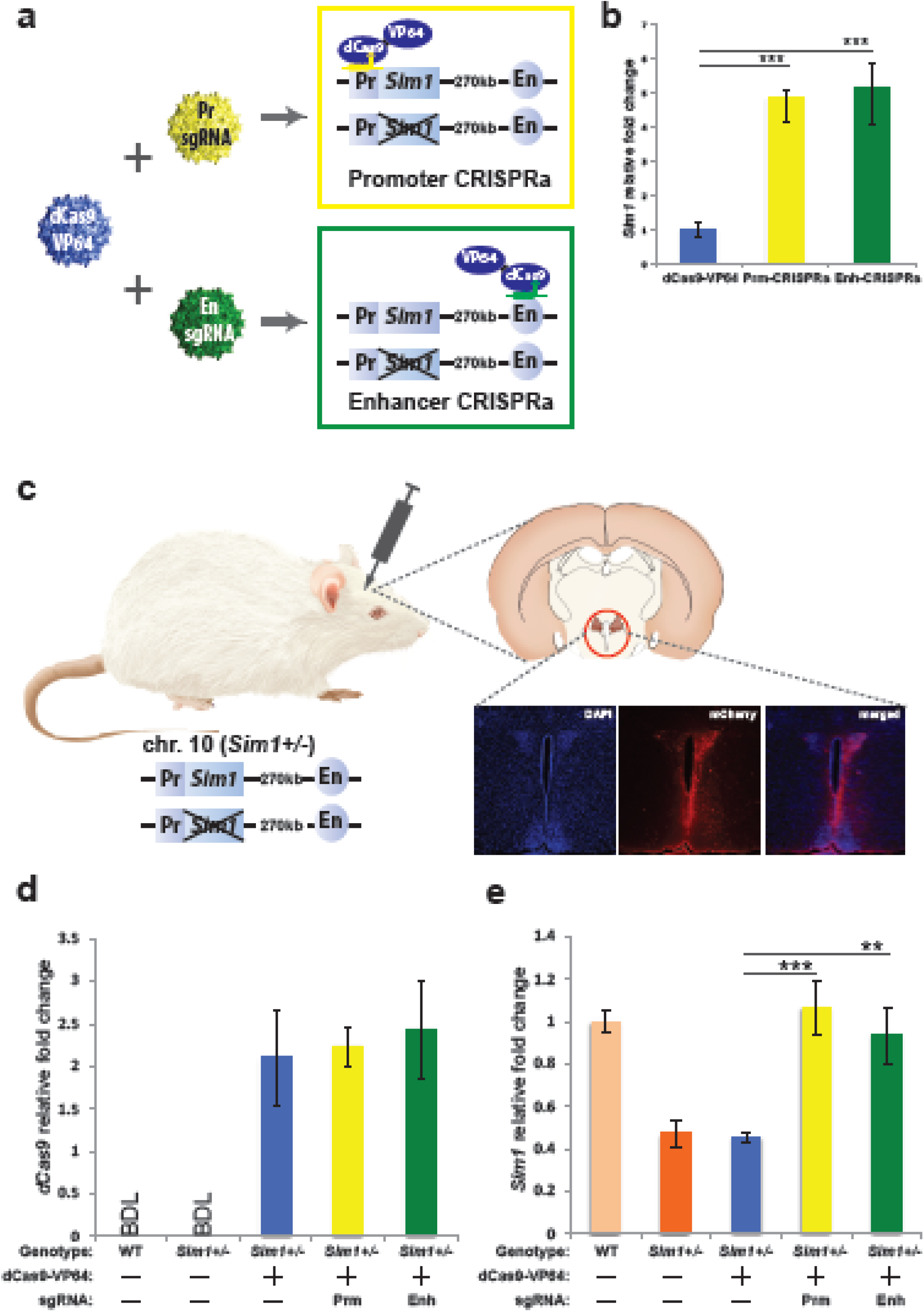
CRISPRa *Sim1* overexpression *in vitro* and *in vivo* using AAV. **a**, Schema showing the various AAVs used for *Sim1* CRISPRa. **b**, AAV CRISPRa in Neuro-2A cells using virons containing: *pCMV-dCas9-VP64* (dCas9-VP64), *pCMV-dCas9-VP64* along with *pSim1Pr-mCherry* (Prm-CRISPRa) and *pCMV-dCas9-VP64* along with *pSCE2En-mCherry* (Enh-CRISPRa). Results are expressed as mRNA fold-increase normalized to beta-actin using the ΔΔCT method. The mean values±s.d. were obtained from 3 independent experiments. *** = p-value < 0.0005 (ANOVA, Tukey test). **c**, Schema showing the stereotactic injection location in the PVN (red circle) followed by immunohistochemistry results from *pCMV-dCas9-VP64* + *pSim1Pr-mCherry* injected hypothalamus of 20 week old mice showing DAPI staining, mCherry expression and both of them merged. **d-e**, *dCas9* (**d**) and *Sim1* (**e**) mRNA expression from non-injected wild-type and *Sim1*^+/−^ mice along with *pCMV-dCas9-VP64* (dCas9-VP64), *pCMV-dCas9-VP64* + *pSim1Pr-mCherry* (Prm-CRISPRa) and *pCMV-dCas9-VP64* + *pSCE2En-mCherry* (Enh-CRISPRa) injected *Sim1*+/− mice. Three mice were used for each genotype. The mean values±s.d were determined based on mRNA fold-increase compared to wild-type mice and normalized to beta-actin using the ΔΔCT method for *Sim1* expression and relative *beta-actin* ΔCT log2 for *dCas9* expression. BDL = below detectable levels.

Next, we performed stereotactic injections to deliver virus carrying *pCMV-dCas9-VP64* and either *pU6-Sim1Pr-CMV-mCherry* (Prm-CRISPRa-AAV) or *pU6-SCE2-CMV-mCherry* (Enh-CRISPRa-AAV) into the PVN of the hypothalamus of *Sim1*^+/−^ mice at four weeks of age, before they start developing obesity. As an injection-based negative control, we also injected *Sim1*^+/−^ mice with *pCMV-dCas9-VP64* virus only. We tested for the expression of our sgRNA-CMV-mCherry cassette by performing immunostaining on the hypothalamus of injected mice and found it to be expressed in the PVN (**Fig. 4c**). To test whether *Sim1* expression levels were increased by delivering CRISPRa-AAV to the hypothalamus of *Sim1*^+/−^ mice, we measured mRNA expression levels for both *dCas9* and *Sim1* from 11 week old AAV injected mice. *dCas9* was found to be expressed in the hypothalamus of all our *pCMV-dCas9-VP64* AAV injected mice (**Fig. 4d**). *Sim1* upregulation was observed in both Prm-CRISPRa-AAV and Enh-CRISPRa-AAV injected hypothalami, but not in mice injected with only *pCMV-dCas9-VP64-AAV* (**Fig. 4e**). To observe the extent that can be achieved of *Sim1* upregulation, we performed Prm-CRISPRa-AAV injections into the hypothalamus of wild-type mice using two different titers. The highest upregulation level that we observed was up to 1.8 fold with the higher viral titer (**Extended Data Fig. 8**).

We next set out to test whether *Sim1* upregulation via CRISPRa AAV can lead to a reduction in body weight. CRISPRa-AAV injected *Sim1*^+/−^ mice were then measured for body weight up to 11 weeks of age (**Fig. 5a**). We observed a significant weight reduction in the Prm-CRISPRa-AAV or Enh-CRISPRa-AAV injected mice compared to the *Sim1*+/− or *pCMV-dCas9-VP64-AAV* injected *Sim1*^+/−^ mice (**Fig5. b**). While many of the injected mice were analyzed in the aforementioned gene expression studies, a few were maintained and showed significant weight reduction compared to the *Sim1*^+/−^ or *pCMV-dCas9-VP64-AAV* injected *Sim1*^+/−^ mice nine months post injection (**Fig. 5c-d**). These results show that CRISPRa-AAV-mediated upregulation could be developed as a potential gene therapy tool to treat haploinsufficiency.

**Figure 5.**
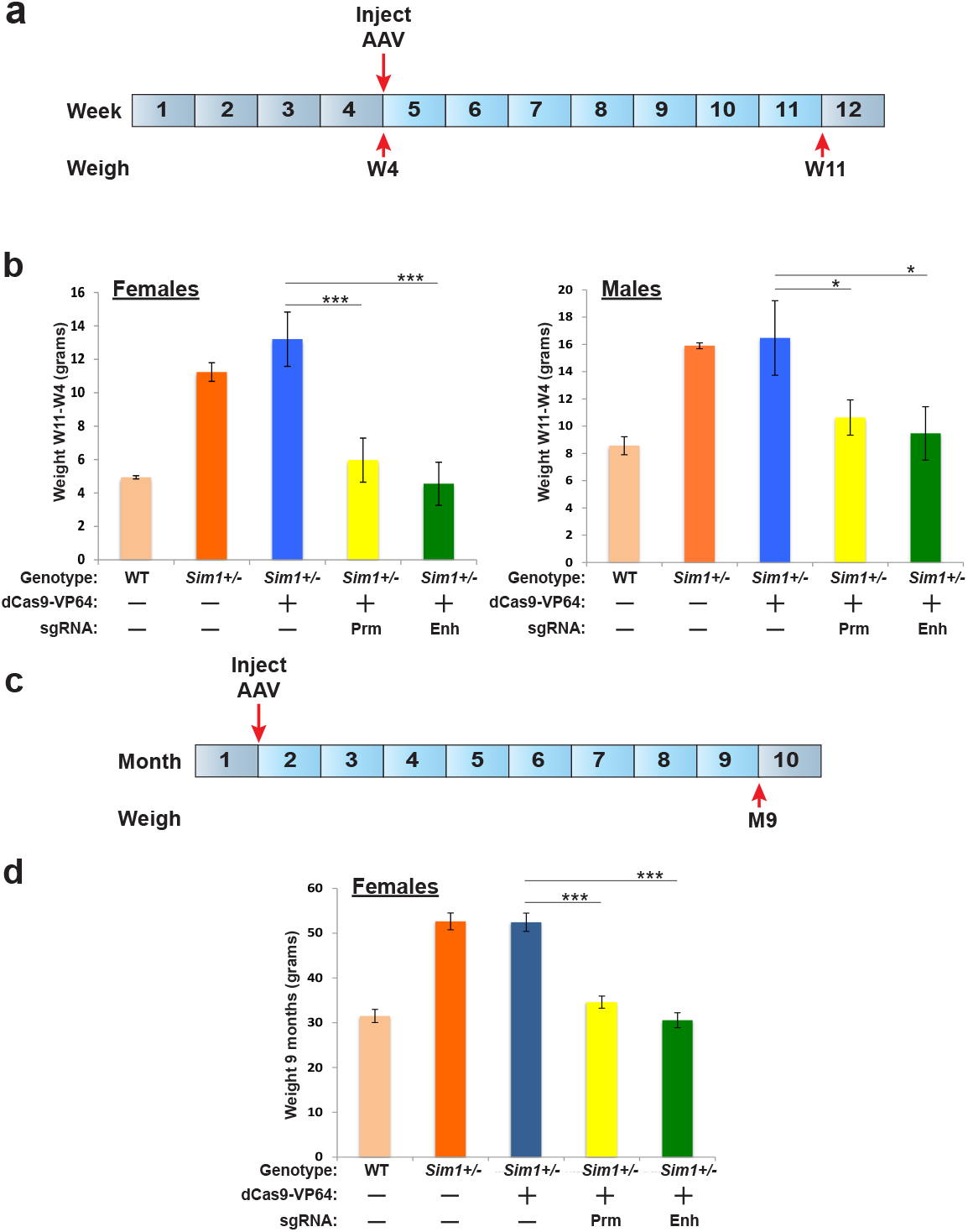
CRISPRa-AAV injection in PVN decreases weight gain in *Sim1*^+/−^ mice. **a**, Timeline for weight measurement post CRISPRa-AAV injection in PVN. **b**, Weight gain determined over a 7 week period from *Sim1+/−* mice injected with *pCMV-dCas9-VP64* (dCas9-VP64), *pCMV-dCas9-VP64* + *pSim1Pr-mCherry* (Prm-CRIPSRa) *pCMV-dCas9-VP64* + *pSCE2En-mCherry* (Enh-CRISPRa) compared to un-injected wild-type littermates and *Sim1*^+/−^ mice. Mean values±s.d are shown from at least 3 mice per gender and genotype. * = p-value < 0.001 *** = p-value < 0.0005 n.s = non-significant; (ANOVA, Tukey test). **c**, Monthly timeline for weight measurement post CRISPRa-AAV injection in PVN. **d**, dCas9-VP64, Prm-CRIPSRa and Enh-CRISPRa compared to un-injected wild-type littermates and *Sim1*^+/−^ mice nine months post injection. Mean values±s.d are shown from 3 mice per genotype. ***= p-value< 0.0005.

## Discussion

CRISPR-based gene editing is a promising therapeutic technology to correct genetic mutations. However, it currently is not an ideal technology for haploinsufficiency, limited by low non-homologous end joining (NHEJ) efficiencies (*i.e*. editing only a small portion of cells) and the need to custom tailor specific guides and donor sequences for each individual mutation. In addition, it is not a feasible therapeutic strategy for micro-deletions, over 200 of which are known to cause human disease^29^, primarily due to haploinsufficiency. In this study, we used a novel approach to tackle these hurdles and show how a haploinsufficient disease could be corrected by increasing the transcriptional output from the existing functional allele via CRISPRa.

Using CRISPRa targeting for either the promoter or enhancer of *Sim1*, we were able to rescue the obesity phenotype in a tissue-specific manner in mice that are haploinsufficient for *Sim1*. As this therapeutic approach takes advantage of the existing functional allele, it has several benefits: 1) It overcomes the need to custom tailor CRISPR gene editing approaches for different haploinsufficient causing mutations in the same gene. 2) This approach could potentially be used to target two or more genes. As such, it could pose as a potential therapeutic strategy for microdeletions related-diseases that are caused by the heterozygous LoF of more than one gene. 3) CRISPRa-AAV could be used to rescue haploinsufficient diseases caused by genes that are longer than its optimal packaging capability. 4) Tissue-specificity is a major concern for gene therapy. CRISPR-based therapies can take advantage of cis-regulatory elements to guide tissue-specificity (**Fig. 6a**). The availability of large-scale tissue-specific maps of gene regulatory elements could provide ample candidates to use for this therapeutic approach. We observed distinct difference in tissue specific activation of *Sim1* in our study based on the targeted *cis*-regulatory element, which can be attributed to chromatin accessibility of the locus in various tissues. Previous large-scale Cas9 and dCas9 cell culture screens have shown a targeting preference for regions with low nucleosome occupancy^30,31^. Active promoters or enhancers would have lower nucleosome occupancy, thus being more amenable to dCas9 targeting.

**Figure 6.**
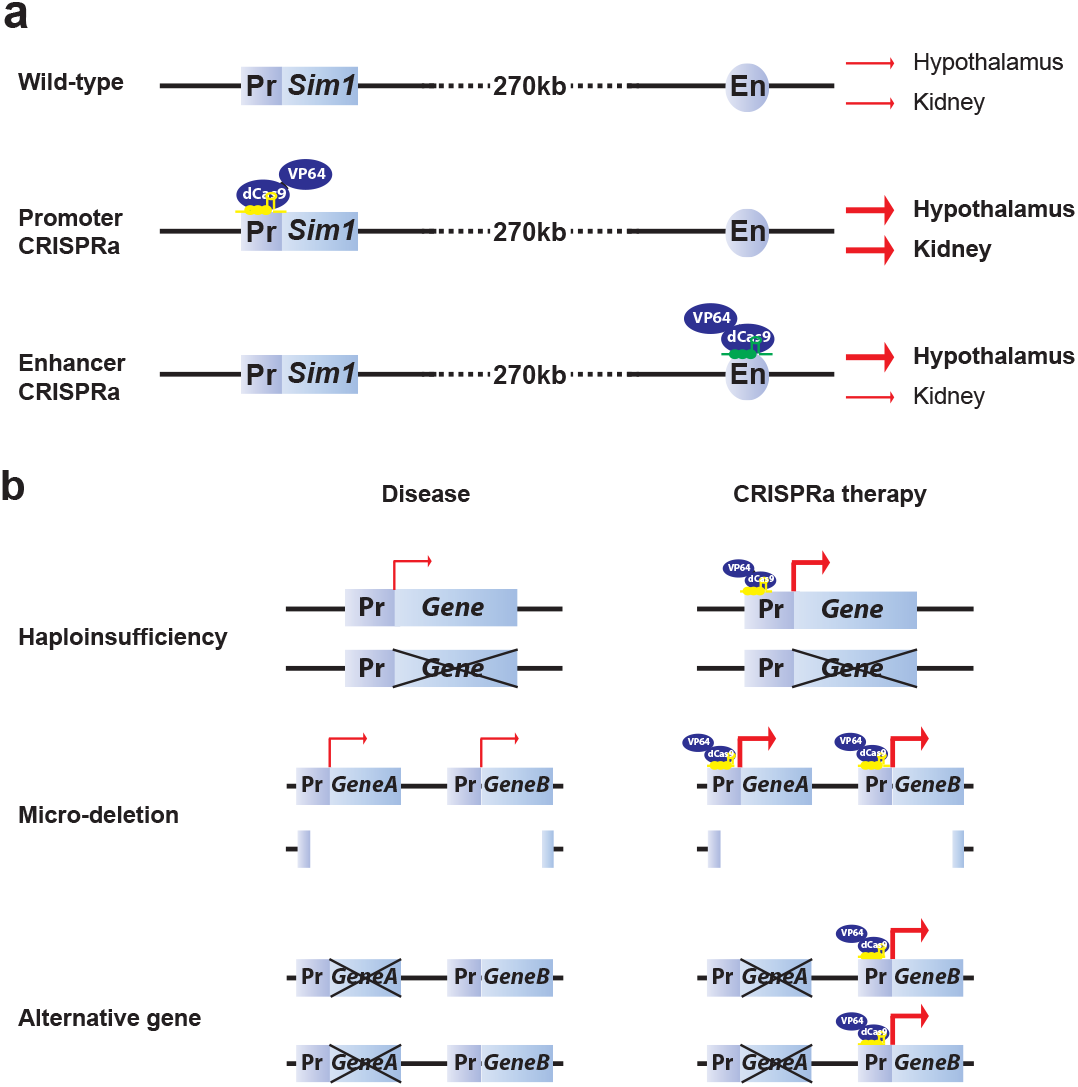
CRISPRa therapeutic potential. **a**, Tissue-specific differences in gene activation due to the type of targeted cis-regulatory element (promoter or enhancer). **b**, CRISPRa can be used as a therapeutic tool to rescue haploinsufficient disease by upregulating the expression of the endogenous functional allele. It can also be used to upregulate a gene or genes that are deleted in micro-deletions or an alternate gene with a similar function to the disease mutated gene.

Our dCas9-VP64 mouse and AAV vectors can be a useful tool for targeted gene activation *in vivo* by delivering sgRNA/s targeted to a specific gene/s in certain tissues/cell types. This approach could be used to assess gene-gene interactions or for the identification of the target gene/s of a specific regulatory element *in vivo* by measuring its expression level following activation. Another potential area of study could be neuronal circuit manipulation. Discrepancies between acute and chronic neuronal circuit manipulations have been observed^32^ which could potentially be addressed by our AAV-CRISPRa and Transgenic-CRISPRa strategies respectively.

Haploinsufficiency of *Sim1* causes obesity both in mice^17^ and humans^13^. Whether this is caused by the reduction in PVN size during development that is observed in *Sim1*^+/−^ mice^17^ or by disturbed energy homeostasis during adulthood was an area of major research. The obesity phenotype observed in the postnatal conditional knockout of hypothalamic Sim1^18^, reinforced the hypothesis that *Sim1* does indeed have a role in energy homeostasis later during adulthood. The observed weight reduction in wild-type *Sim1* CRISPRa mice, the upregulation of *Mc4r* and *Oxt* in Prm-CRISPRa and Enh-CRISPRa mice and the ability to rescue the obesity phenotype via CRISPRa AAV injections into the hypothalamus of 4 week old mice, further corroborates this role. Abrogation of *Mc4r* signaling is the hallmark of most polygenic and monogeneic obesity phenotypes. Conditional postnatal deficiency of *Sim1* leads to reduced levels of Mc4r signaling. As *Sim1* was shown to be an integral downstream component of the leptin-Mc4r pathway^18^, future testing of whether *Sim1* CRISPRa targeting could provide a potential therapy for conditions that disrupt the leptin signaling pathway will be of extreme interest.

Despite technological advances in CRISPR-based therapeutic intervention, our understanding of the long-term side effects of CRISPR expression and its off-targeting effects *in vivo* still remains largely unknown, which also holds true for our current study. Anti-CRISPR genes^33^ or conditional activation or silencing of our CRISPRa system could be able to address these concerns in the future. Furthermore, there is also a need to develop CRISPRa/i tools to modulate gene dosage, so as to be able to optimize transcriptional output for certain diseases where higher or lower activation levels might be needed. In this study, we used VP64 as our activator, due to its known weak activation capacity^23^, which fits with our need to obtain levels of gene expression that are similar to having two normal alleles. CRISPRa based gene activation is highly dependent upon the nature of the fused activator^23^, sgRNA target^30^ and would need to be optimized along with the delivery method. In addition, the use of a shorter Cas9, such as the *Staphylococcus aureus^34^* or *Campylobacter jejuni^35^*, could increase viral titers. Similarly, AAV has its own share of caveats such as inflammation and immunological reactions^36,37^ or association with hepatocellular carcinomas^38–40^ Further development of AAV therapeutic tools could overcome several of these limitations.

To our knowledge, this is the first study to use CRISPRa as a therapeutic tool in a mouse disease model. We demonstrate that CRISPRa can be used to activate genes not only by targeting their promoters, but by also targeting distal cis-regulatory elements such as enhancers. Previous studies have shown that these elements can be viable therapeutic targets. For example, by targeting a globin enhancer with zinc finger nucleases fused to a chromatin looping factor, the LIM domain binding 1 (*LDB1*) gene, activation of fetal hemoglobin was achieved *in vitro*, providing a potential therapy for sickle cell disease^41^. In another study, re-activation of fetal hemoglobin was achieved by deactivating the enhancer of its repressor B-cell CLL/lymphoma 11A (*BCL11A*) using CRISPR gene editing^42^. Our study provides a novel approach that also takes advantage of cis-regulatory elements for therapeutic purposes. There are numerous diseases that are caused by lower gene dosage that could potentially be treated with CRISPRa therapy (**Fig. 6b**). In addition, several human diseases could potentially be rescued by the activation of another gene with a similar function (**Fig. 6b**). These could include for example *Utrophin* for Duchenne Muscular Dystrophy^43^, survival of motor neuron 2 (*SMA2*) for Spinal Muscular Atrophy (SMA; ^44^) or the aforementioned fetal globin for sickle cell disease. Further development of this technology could provide a potential therapy for patients inflicted with these diseases.

## Methods

### Plasmids

The *pMSCV-LTR-dCas9-VP64-BFP* vector, encoding a mammalian codon-optimized *Streptococcus pyo9enes* dCas9 fused to two C-terminal SV40 NLSs and tagBFP along with a VP64 domain and the U6-sgRNA-CMV-mCherry-T2A-Puro plasmids were used for cell line transfections (both kind gifts from Drs. Jonathan S. Weissman and Stanley Qi). sgRNAs (**Supplementary Table 1**) were cloned using the In-Fusion HD-cloning kit (Clontech) following the manufacturer’s protocol into the *BstXI* and *XhoI* sites. Mouse knockin vectors were generated by cloning dCas9-VP64 and U6-sgRNA-CMV-mCherry expression cassettes from the aforementioned vectors into the TARGATT (CAG + Poly A) plasmid (Applied StemCell). For AAV vectors, *pcDNA-dCas9-VP64* (Addgene 47107), and U6-sgRNA-CMV-mCherry-WPREpA were cloned replacing the *Ef1a-FAS-hChR2(H134R)-mCherry-WPRE-pA* with that of our U6-sgRNA-CMV-mCherry-WPREpA into the backbone of *pAAV-Ef1a-FAS-hChR2(H134R)-mCherry-WPRE-pA* (Addgene 37090).

### AAV production

AAV DJ serotype particles were produced using the Stanford Gene Vector and Virus core. The packaging load for *pCMV-dCas9-VP64* was 5.4kb and for *pU6-Sim1Pr-CMV-mCherry* and *pU6-SCE2-CMV-mCherry* 2.5kb. Genomic titers were ascertained by WPRE and ITR probes to be 1.40E^10^ viral genome (vg)/ml for *pCMV-dCas9-VP64* and around 3.30E^13^ vg/ml for *pU6-Sim1Pr-CMV-mCherry* and 2.20 E^13^ vg/ml for *pU6-SCE2-CMV-mCherry*.

### Cell culture

Neuroblastoma 2a cells (Neuro-2a; ATCC^®^ CCL-131) were grown following ATCC guidelines. Plasmids were transfected into Neuro-2a cells using X-tremeGENE HP DNA transfection reagent (Roche) following the manufacturer’s protocol. AAV particles were infected into Neuro-2a cells at different multiplicity of infection (MOIs; **Extended Data Fig. 9**). Neuro2a cells were harvested 48 hours post transfection and 5 days post infection to isolate RNA for qRT-PCR analysis.

### RNA isolation quantitative reverse-transcription PCR

RNA was isolated from cells or tissues using RNeasy Mini Kit (Qiagen) following the manufacturer’s protocol. For mice, animals were euthanized and tissues were harvested directly into the RNA lysis buffer of the RNeasy Mini Kit. The hypothalamus was dissected using a mouse Brain Matrix and slicers from Zivic Instruments. cDNA was prepared using SuperScript III First-Strand Synthesis System (Invitrogen) using the manufacturer’s protocol along with DNaseI digestion. qPCR was performed using SsoFast EvaGreen Supermix (Biorad) using primers in **Supplementary Table 1**. The results were expressed as fold-increase mRNA expression of the gene of interest normalized to either beta-actin, *Rpl38* or *Elf3* expression by the ΔΔCT method followed by ANOVA and Tukey test for statistical analysis. Reported values are the mean and standard error of the mean from three independent experiments performed on different days (N=3) with technical duplicates that were averaged for each experiment.

### Chromatin immunoprecipitation

Fresh tissue was homogenized using a hand-held dounce homogenizer, cross-linked in Phosphate buffer saline (PBS) containing 1% formaldehyde for 10 minutes, quenched with 125mM Glycine for 5 minutes and washed three times with PBS. Cross-linked tissue pellet was processed further for chromatin immuoprecipitation using ‘Low cell Chip Kit’ Diagenode (Catalog number C01010072) following the manufacturer’s protocol. Cas9 polyclonal antibody from Diagenode (Catalog number C15310258) was used for the pull down. Enrichment of target regions was assessed by RT-qPCR using SsoFast EvaGreen Supermix (Bio-Rad) and primers listed in **Supplementary Table 1**. Results were expressed as %input using the ΔCT method. Reported values are the mean and standard error of the mean from three independent experiments performed on different days (N=2) with technical duplicates that were averaged for each experiment.

### Mice

*Sim1*^+/−^ mice^17^ on a mixed genetic background were obtained as a kind gift from Dr. Jacques Michaud’s lab. In these mice, a 1 kb fragment containing 750bp of the 5’ region, the initiation codon, and the sequence coding for the basic domain (the first 17 amino acids) was replaced by a *Pgk–neo* cassette, that was used for genotyping (see **Supplementary Table 1** for primers) using KAPA mouse genotyping kit (KAPA Biosystems). To generate dCas9-VP64 and sgRNA mice we used TARGATT technology^24^. DNA for injection was prepared and purified as mini-circles using the TARGATT Transgenic Kit, V6 (Applied StemCell). The injection mix contained 3 ng/μL DNA and 48 ng/μL of *in vitro* transcribed φC31o mRNA in microinjection TE buffer (0.1 mM EDTA, 10 mM Tris, pH 7.5) and injections were done using standard mouse transgenic protocols^45^. dCas9-VP64 was inserted into the mouse *Hipp11* locus and sgRNAs into the *Rosa26* locus. Mice were genotyped using the KAPA mouse genotyping kit. F0 H11P TARGATT knockins were assessed using PCR primers SH176 + SH178 + PR432 and for ROSA26 primers ROSA10 + ROSA11 + PR432 described in^24^ along with vector insertion specific dCas9-VP64 primers as well as mCherry specific primers (described in **Supplementary Table 1**). All mice were fed *ad libitum* Picolab mouse diet 20, 5058 containing 20% protein, 9% fat, 4% fiber for whole study. Calories provided by: protein 23.210%, fat (ether extract) 21.559% and carbohydrates 55.231%. All animal work was approved by the UCSF Institutional Animal Care and Use Committee.

### Mouse body weight measurements

*H11P^CAG-dCas9-VP64^*, *ROSA26^Sim1Pr-sgRNA^* and *ROSA26^SCE2En-sgRNA^* mice were mated with FVB mice for 3-5 generations to assess germline transmission. Three independent integrants were used from each line to set up matings. *H11P^CAG-dCas9-VP64^* were mated with *Sim1*^+/−^ and subsequent *Sim1*^+/−^ X *H11P^CAG-dCas9-VP64^* mice were crossed with either *ROSA26^Sm1Pr-sgRNA^* or *ROSA26^SCE2En-sgRNA^* to generate mice having all three unlinked alleles. Mice were maintained at Picodiet 5058 throughout the study and at least 10 females and 10 males from all genotypes (wild-type, *Sim1*^+/−^, *Sim1*^+/−^ X *H11P^CAG-dCas9-VP64^*, *Sim1*^+/−^ X *H11P^CAG-dCas9-VP64^* X *ROSA26^Sim1Pr-sgRNA^*, *Sim1*^+/−^ X *H11P^CAG-dCas9-VP64^* X *ROSA26^SCE2En-sgRNA^*) were measured for their body weights from 4-16 weeks of age on a weekly basis.

### Mouse metabolic profiling

Metabolic rates from individual mice were measured using the Columbus Instruments Comprehensive Lab Animal Monitoring System (CLAMS; Columbus Instruments). Mice were single housed and acclimatized on powdered picodiet 5058 for 3-4 days before performing the metabolic monitoring. We individually housed mice in CLAMS units and measurements were carried out over 4-5 days. The temperature was maintained at 22°C and oxygen and carbon dioxide were calibrated with ‘Air reference’ set at 20.901 and 0.0049. Three males and three females from each genotype: wild-type littermates, *Sim1*^+/−^, *Sim1*^+/−^ X *H11P^CAG^’^dCas9^’^VP64^* X *ROSA26^Sim1Pr-sgRNA^*, *Sim1*^+/−^ X *H11P^CAG-dCas9-VP64^* X *ROSA26^SCE2En-sgRNA^* were measured for the following metabolic parameters: VCO2, VO2, RER, food intake and activity monitoring. Metabolic data was analyzed using CLAX support software (Columbus Instruments).

### Body composition analysis

Body composition was measured using either Dual Energy X-ray Absorptiometry (DEXA) or Echo Magnetic Resonance Imaging (EchoMRI; Echo Medical System). For DEXA, mice were anesthetized using isoflurane and measured for bone mineral density and tissue composition (fat mass and lean mass) using the Lunar PIXImus. EchoMRI (Echo Medical System) was used to measure whole body composition parameters such as total body fat, lean mass, body fluids, and total body water in live mice without the need for anesthesia or sedation.

### Stereotaxic injections

Four week-old *Sim1*^+/−^ males or females, weighing between 22 and 26 grams, were housed individually in cages for at least 2 days before surgical interventions. Mice were anesthetized with a 100 mg/kg Avertin intraperitoneal injection. The skull was immobilized in a stereotaxic apparatus (David Kopf Instruments). The stereotaxic coordinates for injection into the PVN were 0.80 mm caudal to bregma, 0 mm at the midline, and 5.2 mm below the surface of the skull. A 1.5 mm hole was created in the cranium by circular movements using a hand-held Dumont 5-45 tweezers (Fine Science Tools). Using a 31 gauge 1ul Hamilton microsyringe, we injected a dose of 0.5X10^7^ vg/ml of sgRNA-AAV along with 2.5X10^6^ vg/kg of dCas-VP64-AAV, in a total injection volume of 1ul per animal into the PVN unilaterally over a 10 minute period. This titer and double the amount (1X10^7^ vg/ml of sgRNA-AAV along with 5X10^6^ vg/kg of dCas-VP64-AAV) were also injected into 5 week old FVB mice (**Extended Data Fig. 7**). After AAV delivery, the needle was left in place for 20 minutes to prevent reflux and slowly withdrawn in several steps, over 10 minutes. Mice were administered two doses of buprenorphine (100mg/kg) before and 24 hours post-surgery. Immunostaining for mCherry, as described below, was used to validate PVN injection coordinates 2-12 weeks following injection in several mice. Mice were maintained on a picodiet 5058 and weighed on a weekly basis.

### Immunostaining

For immunostaining, mice were anesthetized with pentobarbital (7.5 mg/0.15 ml, i.p.) and transcardially perfused with 10ml of heparinized saline (10 U/ml, 2 ml/min) followed by 10ml of phosphate-buffered 4% paraformaldehyde (PFA). Brains were removed, postfixed for 24 hours in 4% PFA, and then equilibrated in 30% sucrose in PBS for 72 hours. Brains were coronally sectioned (35 microns for immunostaining, 50 microns for stereology) on a sliding microtome (Leica SM 2000R). Immunohistochemistry was performed as previously described^19,46,47^. Coronal brain sections that had been stored in PBS at 4°C were permeabilized and blocked in 3% normal goat serum/0.3% Triton X-100 for 1 hour and incubated at 4°C overnight using an mCherry antibody at a dilution of 1:500 (Abcam ab167453). Sections were placed in 4,6- diamidino-2- phenylindole (DAPI) (0.2 g /ml; 236276; Roche) for 10 minutes and then mounted on plus coated slides and coverslipped using Vectashield (H-1000; Vector Laboratories). Images of sections containing PVN were captured on a Zeiss Apotome.

## Extended data figure legends

**Extended Data Figure 1 | Coding sequence size distribution of genes associated with haploinsufficiency in humans**. The coding sequence (CDS) size distribution in base pairs (bp) of 3,230 gene loci, estimated to be to intolerant for heterozygous loss of function^3^ (blue) and 300 genes that are known to cause human disease when haploinsufficient^1^ (orange). CDSs longer than 3,500 bp would be difficult to use in conventional AAV-based gene therapy due to packaging size limitations.

**Extended Data Figure 2 | CRISPRa upregulation of *Sim1* in mouse Neuroblastoma-2A cells. a**, Schema of the mouse *Sim1* locus and CRISPRa targeting using two different sgRNAs per target (promoter or SCE2). b, CRISPRa in Neuro-2A cells using either of the two sgRNAs targeting the *Sim1* promoter (Prm-CRISPRa) or enhancer (Enh-CRISPRa). sgRNA-1-Prm, and sgRNA-1-Enh were selected for further studies. Results are expressed as mRNA fold-increase normalized to *beta-actin* using the ΔΔCT method. The mean values±s.d. were obtained from 3 independent experiments. **** = p-value < 0.0001 (ANOVA, Tukey test).

**Extended Data Figure 3 | Generation of mouse transgenics and mating scheme. a**, Diagram showing the knockin strategy to generate CRISPRa transgenics. Transgenes were inserted into either the *Hipp11* or *Rosa26* locus having three attP sites by pronuclear microinjection of minicircles carrying the transgene along with Phi-integrase mRNA. **b**, Mating scheme to combine the three unlinked loci. *Sim1*^+/−^ mice were crossed to *H11P^CAG-dCas9-VP64^* and mice having both alleles were then crossed to either *ROSA26^Sim1Pr-sgRNA^* or *ROSA26^SCE2En-sgRNA^* mice to generate Prm-CRISPRa and Enh-CRISPRa mice respectively.

**Extended Data Figure 4 | Weekly weight measurements for the various mouse genotypes**. Weekly weight measurements of wild-type, *Sim1*^+/−^, *H11P^CAG-dCas9-VP64^* (dCas9-VP64), *ROSA26^Sim1Pr-sgRNA^*, *ROSA26^SCE2En-sgRNA^*, *Sim1*^+/−^ X *H11P^CAG-dCas9-VP64^*, *Sim1*^+/−^ X *ROSA26^Sim1Pr-sgRNA^* and *Sim1*^+/−^ X *ROSA26^SCE2En-sgRNA^*. At least 6 male and female mice were measured per genotype (number of mice, n, is shown next to each genotype). Mean values±s.d are shown.

**Extended Data Figure 5 | Mouse physical activity measurements**. Physical activity measurements of wild-type, *Sim1*^+/−^, *H11P^CAG-dCas9-VP64^*X *ROSA26^Sim1Pr-sgRNA^* (Prm-CRISPRa) and *H11P^CAG-dCas9-VP64^*X *ROSA26^SCE2En-sgRNA^* (Enh-CRISPRa) mice using the CLAMS system. Three males and three female mice were used per genotype.

**Extended Data Figure 6 | *Sim1* mRNA expression in multiple tissues**. *Sim1* mRNA expression for the following genotypes: wild-type, *Sim1*^+/−^ and *H11P^CAG-dCas9-VP64^* X *ROSA26^Sim1Pr-sgRNA^* (Prm-CRISPRa). For the hypothalamus, kidney, lung, liver data was taken from 4 mice (2 females and 2 males) and for the trachea, skeletal muscle, upper spinal cord and visceral adipose tissue (V. adipose tissue) data was taken from 2 mice. The mean values±s.d for all experiments were determined based on mRNA fold-increase compared to wild-type and normalized to *beta-actin* or *Rpl38* using the ΔCT method.

**Extended Data Figure 7 | mRNA and ChIP-qPCR analyses. a-b**, *Ascc3* (**a**) and *Gprc6a* (**b**) mRNA expression levels in the hypothalamus for the following genotypes: wild-type, *Sim1+/−*, Prm-CRISPRa and Enh-CRISPRa from 4 mice (2 females and 2 males). Results are expressed as mRNA fold-increase normalized to *beta-actin* using the ΔΔCT method. The mean values±s.d. were obtained from 3 independent experiments. **c-d**, ChIP-qPCR using two *H11P^CAG-dCas9-VP64^* X *ROSA26^Sim1Pr-sgRNA^* (Prm-CRISPRa) mice (**c**) and two *H11P^CAG-dCas9-VP64^*X *ROSA26^SCE2En-sgRNA^* (Enh-CRISPRa) mice (**d**) for the *Ascc3* and *Gprc6a* promoters, a negative control region (*Mc4r* promoter) and either the *Sim1* promoter (**c**) or *SCE2* (**d**). Results are expressed using the ΔCT% input method. The mean values±s.d. were obtained from two independent experiments.

Extended Data Figure 8 | Prm-CRISPRa-AAV injections in wild-type mice. *Sim1* mRNA expression levels from 5 week old FVB mice injected with *Prm-CRISPRa-AAV* using a dose of 0.5X10^7^ vg/ml of sgRNA-AAV along with 2.5X10^6^ vg/kg of dCas-VP64-AAV (1X titer), a dose of 1X10^7^ vg/ml of sgRNA-AAV along with 5X10^6^ vg/kg of dCas-VP64-AAV (2X titer) and *pCMV-dCas9-VP64* (dCas9-VP64) compared to un-injected mice. Measurements were taken 3 weeks post injection and are expressed as fold-increase compared to un-injected mice. The mean values±s.d. were obtained from four different mice for each condition and three independent experiments.

Extended Data Figure 9 | CRISPRa upregulation of *Sim1* in Neuro-2a cells using different AAV MOIs. AAV CRISPRa in Neuro-2A cells using virons containing: *pCMV-dCas9-VP64* (dCas9-VP64), *pCMV-dCas9-VP64* along with *pSim1Pr-mCherry* (Prm-CRISPRa) and *pCMV-dCas9-VP64* along with *pSCE2En-mCherry* (Enh-CRISPRa). Two different multiplicity of infection (MOI) were used: 5,000 and 7,500 viral genome (vg)/ml. Results are expressed as mRNA fold-increase normalized to *beta-actin* using the ΔΔCT method. The mean values±s.d. were obtained from 3 independent experiments.

**Supplementary Table 1** | sgRNA and primer sequences. For the sgRNAs, in bold are the sequences used for mice and AAV.

## Acknowledgments

We want to thank Dr. Jacques Michaud for the *Sim1*^+/−^ mice, Dr. Mee J. Kim for the SCE2 hypothalamus picture, Drs. Jonathan S. Weissman and Stanley Qi for the dCas9-VP64 plasmids, Dr. Christophe Paillart for all the assistance with the metabolic profiling, Sunpreet Kaur Matharu for graphic assistance with the figures and Drs. Michael T. McManus, Patrick Devine, Walter L. Eckalbar and Sajad H. Ahanger for helpful discussions. This article was supported in part by a grant from the National Institute of Diabetes and Digestive and Kidney Diseases (NIDDK) 1R01DK090382, the UCSF Nutrition Obesity Research Center funded by that National Institute of Health P30DK098722 and the UCSF School of Pharmacy 2017 Mary Anne Koda-Kimble Seed Award for Innovation. N.A. is also supported by grants by the National Human Genome Research Institute (NHGRI) and Division of Cancer Prevention, National Cancer Institute grant number 1R01CA197139, National Institute of Mental Health grant number 1R01MH109907, National Institute of Child & Human Development 1P01HD084387 and NHGRI grant number 1UM1HG009408. S.R. was supported by the Royal Golden Jubilee Ph.D. Program grant number PHD/0071/2554. C.V. is also supported by NIDDK grant numbers R01DK106404 and R01 DK60540-09 and Y.W. from the American Diabetes Association Mentor Based Award-7-12-MN-79. A.H. is supported by the National Institute of General Medical Sciences IRACDA award K12GM081266.

## Author contributions

N.M. and N.A. conceived and designed the study. N.M. and L.M. carried out the cloning and *in-vitro* studies, N.M. and S.R. performed the mouse experiments. N.M. and Y.W. carried out the immunostaining and N.M. preformed the stereotactic surgeries and metabolic profiling. A.H. helped in making Extended Data Fig. 1. N.M., S.R. and N.A. analyzed the data, C.V. and N.A. provided resources and critical suggestions and N.M. and N.A. wrote the manuscript.

